# There is no evidence of a universal genetic boundary among microbial species

**DOI:** 10.1101/2020.07.27.223511

**Authors:** Connor S. Murray, Yingnan Gao, Martin Wu

**Affiliations:** Department of Biology, University of Virginia, 485 McCormick Road, Charlottesville, VA 22904

## Abstract

A fundamental question in studying microbial diversity is whether there is a species boundary and if the boundary can be delineated by a universal genetic discontinuity. To address this question, Jain et al. computed the pairwise average nucleotide identity (ANI) of 91,761 microbial (bacterial and archaeal) genomes (the 90K genome dataset) and found that the ANI values from the 8 billion comparisons follow a strong bimodal distribution^1^. The authors concluded that a clear genetic discontinuum and species boundary were evident from the unprecedented large-scale ANI analysis. As a result, the researchers advocated that an ANI of 95% can be used to accurately demarcate all currently named microbial species. While the FastANI program described in the paper is useful, we argue that the paper’s conclusion of a universal genetic boundary is questionable and resulted from the substantial biased sampling in genome sequencing. We also caution against being overly confident in using 95% ANI for microbial species delineation as the high benchmarks reported in the paper were inflated by using highly redundant genomes.

## Main Text

To demonstrate our point, we first show that biased within-species sampling can generate the bimodal distribution of ANI observed in the original paper even when the speciation rate is constant and the genetic diversity is continuous. We simulated a phylogenetic tree with 3,000 tips and a constant rate of diversification, estimated from a genome tree of 3,000 bacterial genomes that represents the phylogenetic diversity within the 10,616 NCBI RefSeq complete bacterial genomes (the 10K genome dataset). We calculated ANI between tips using a function that accurately captures the relationship between ANI and the branch length in the real data (Supplementary Fig. 1). Because the diversification rate is constant, the branch lengths follow an exponential distribution expected from a Poisson process. As expected, the frequency of ANI declined monotonically when ANI increased (Fig. 1a). However, when only 30 tips (1% of the tips) were sampled with a protocol that emulated within-species sampling bias (each tip has two very closely related genomes sequenced), the ANI distribution became bimodal (Fig. 1b). The right peak increased in prominence when more tips were sampled with bias (data not shown). Our simulations show that limited within-species sampling bias alone can create the bimodal distribution when genetic diversity is continuous and a universal species boundary does not exist.

**Figure 1.**
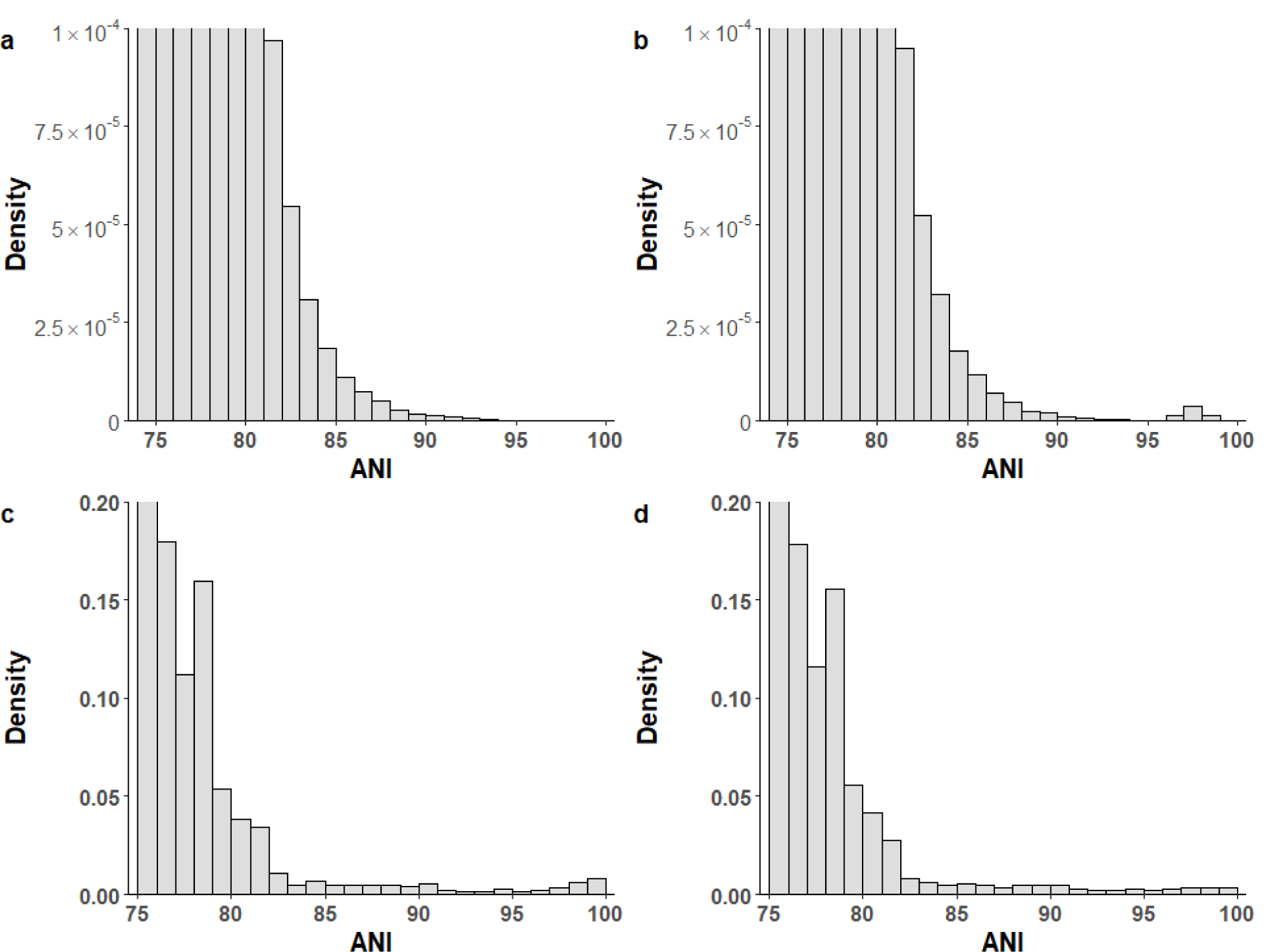
Distribution of ANI values. Top panel: from the phylogenetic simulations a). Comparisons between 3,000 taxa simulated using a phylogenetic tree with 3,000 tips and a constant rate of diversification. b). Comparisons in the same dataset except that 30 of 3,000 taxa each have two very closely related genomes sequenced. Bottom panel: from genomes subsampled from the 90K genome dataset. c). Two genomes were randomly selected from each of the 397 named species with >= 10 genomes. d). Two phylogenetic representative genomes were selected for each of the 397 named species with >= 10 genomes.

In the original 90K genome dataset, 33% of named species have been sequenced at least twice. As cultivation bias is widespread, next we show that there is substantial within-species sampling bias in the genome datasets. For the model organism *Escherichia coli*, its 602 complete genomes in the 10K genome dataset only represent 22% of the diversity captured by the 16S rRNA gene in the GreenGene database (Supplementary Fig. 2). For species with at least 100 genomes, on average the first 75.8% [range: 50.9-97.7%] of dropped genomes contribute less than 5% of the genetic diversity of the species as measured by the branch length (Fig. 2). As demonstrated in our phylogenetic simulation, these highly redundant genomes can create the bimodal distribution even when genetic diversity is continuous. Contrary to the authors’ claim, randomly subsampling (e.g., 5 genomes from the same species) from a biased dataset will not correct for the pre-existing sampling bias. When two genomes were sampled randomly from each of the 397 named species with >= 10 genomes in the 90K dataset, the bimodal distribution was evident (Fig. 1c). However, when we reduced the within-species sampling bias by selecting two phylogenetic representative genomes from the same dataset, the distribution flattened near the end (Fig. 1d), further demonstrating that the bimodal distribution is caused by widespread sampling bias within species. In fact, the within-species sampling bias has increased over time when more strains were sequenced based on their medical and economical relevance and not on their phylogenetic positions (Supplementary Fig. 3), thereby producing the consistent bimodal distributions of ANI over different periods of time as observed in the original study^1^.

**Figure 2.**
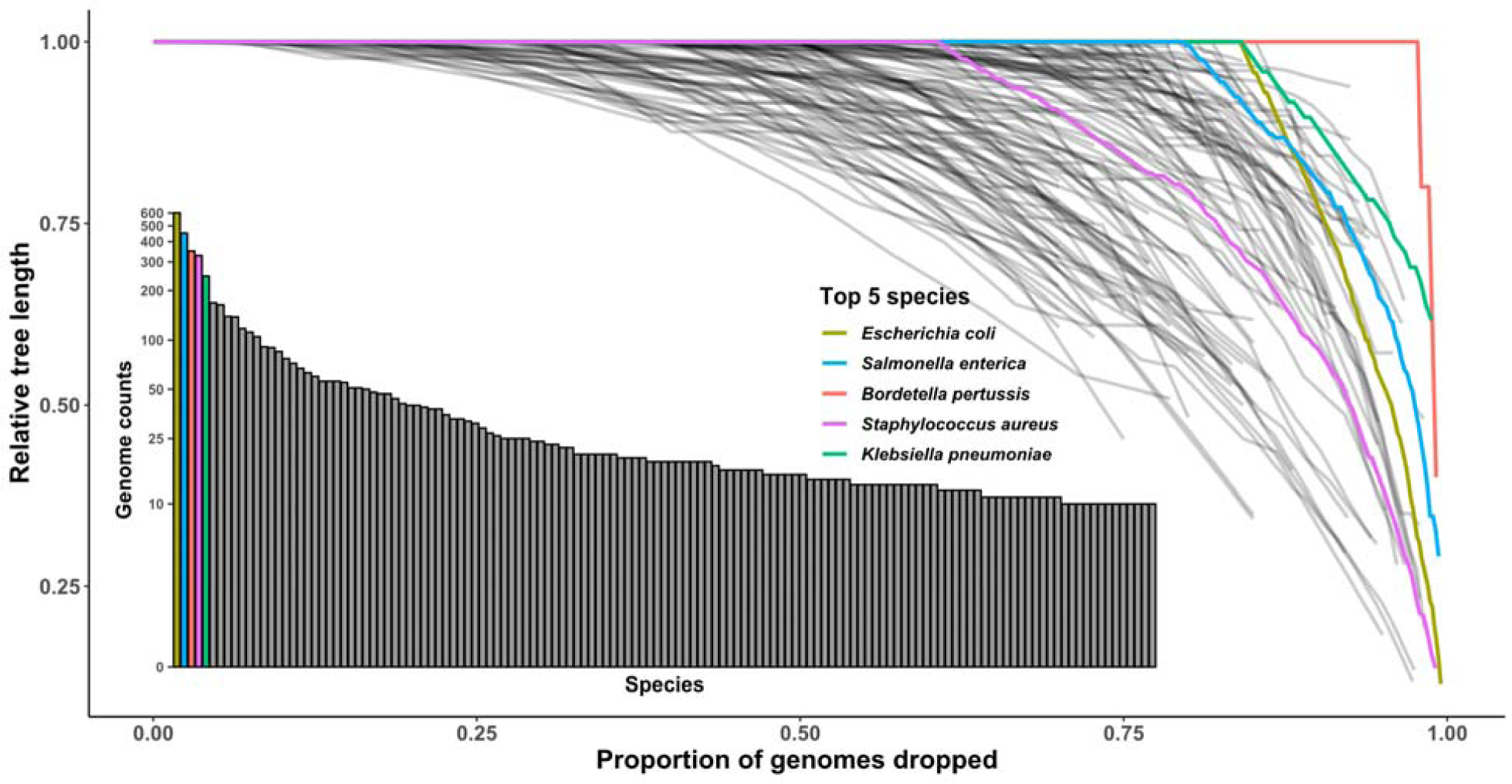
Widespread within-species sampling bias revealed by the Treemmer analysis. Treemmer iteratively drops genomes contributing the least amount of diversity within a phylogenetic tree until only three genomes remain. For each species, the relative tree length (RTL) is plotted against the proportion of genomes dropped at each iteration. Colored lines are the 5 most sequenced species in the 10K genome dataset. Shaded black lines are species with >=10 genomes. The bar chart shows species ranked by the number of genomes sequenced.

The authors indicated that only 0.2% of the 8 billion ANI values are between 83% and 95% and suggested that such a wide gap is evidence of a clear genetic boundary in microbial genomes. In our phylogenetic simulation of 3,000 genomes with continuous genetic diversity, only 0.11% of ANI values span the same region. This is expected because without biased sampling, the vast majority of all pairwise comparisons for a genome will be with distantly related genomes, whose number will always be much larger than the number of closely related taxa. As a result, the fraction of ANI values in the intermediate range [83%-95%] will always be marginal for a decently sized tree (> 50 tips) and will decrease when the number of genomes increases (Supplementary Fig. 4). Our result shows that the low density of ANI values in the gap region does not necessarily indicate a genetic boundary. Instead, it could simply reflect the hierarchical structure of the phylogenetic relationship.

Previous studies have advocated using an ANI cutoff to demarcate bacterial species^2–5^. Based on the unprecedented large-scale ANI analysis of the study, Jain et al. claimed that using the 95% ANI criterion led to both high recall and precision rates (>98.5%) in demarcating microbial species^1^. However, both benchmarks were calibrated with ∼78k named genomes that were highly redundant. For example, there were more than 5,000 *E. coli* genomes in the dataset. As such, the recall and precision rates can be misled by the overly sampled genomes. To better assess the performance of using 95% ANI for species demarcation, we queried the 3,000 representative genomes against the 10K dataset. We found the recall and precision rates were much lower, at 73.4% and 83.3% respectively. The large impact of the extreme sampling bias on the benchmarks is best illustrated by the authors’ own finding that excluding the *E. coli* vs. *Shigella spp*. comparison alone substantially increased their overall precision from 93.1 to 98.7%. Plotting of the intraspecific ANI for intensely sequenced species in the 90K dataset (61 species with >=100 genomes, representing 6 phyla) shows that a universal boundary at 95% ANI clearly does not exist, as ANI values drop well below 95% with few exceptions (Fig. 3).

**Figure 3.**
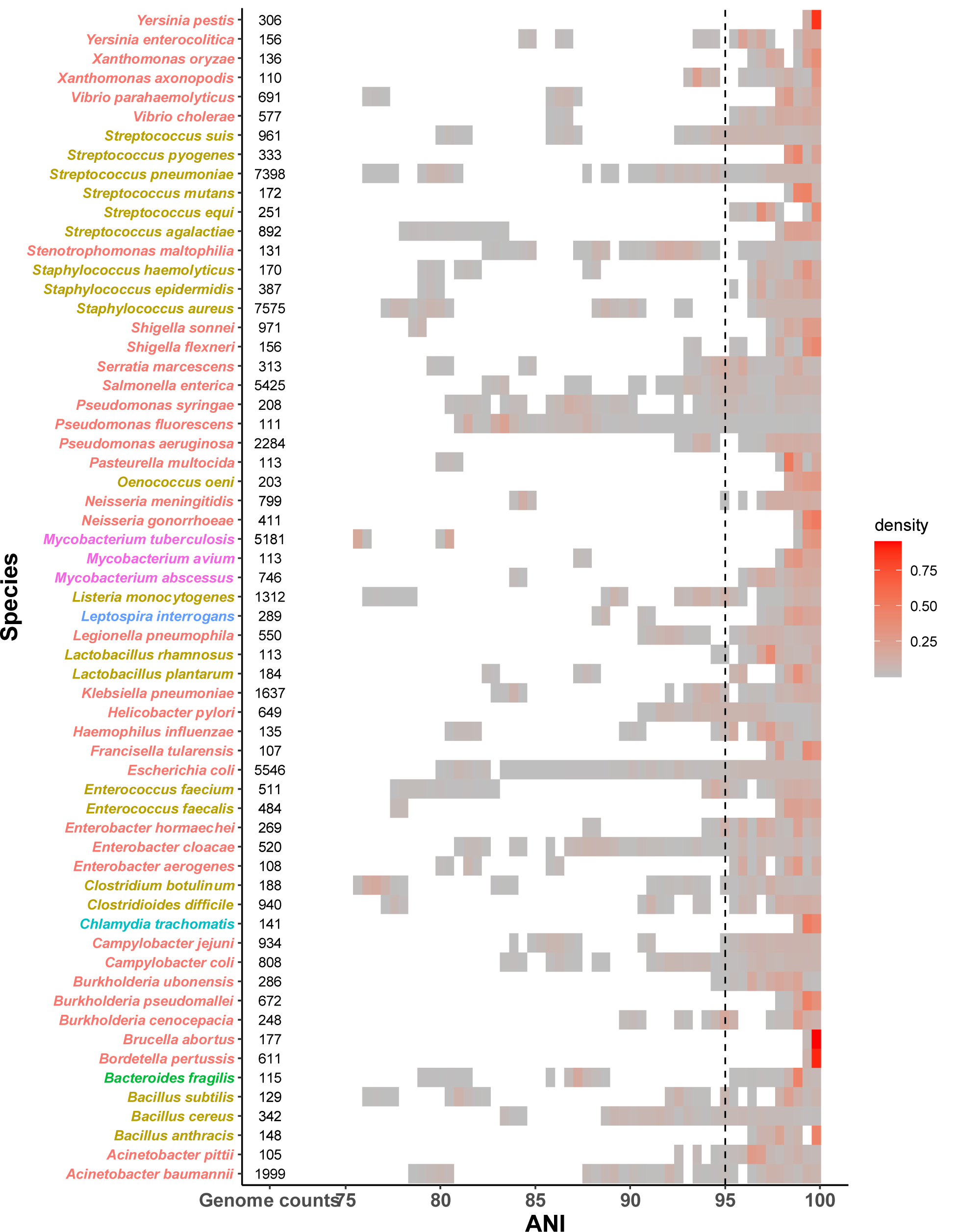
The distribution of ANI for 61 named species with >= 100 genomes in the 90K genome dataset. Each block represents a bin of ANI values between pairs of genomes within the same species and is colored by the ANI density in the bin. Species are colored by phylum. The vertical dotted line represents the 95% ANI threshold. The high ANI density on the right (above the 95% threshold) is due to comparisons between highly redundant genomes from the sampling bias.

Although having no cultivation bias, metagenome assemblies do favor abundant strains over rare ones. This bias can lead to the appearance of genetic clusters when diverse but rare strains in the community are excluded from the assembled genomes. However, a rare strain in one environment can be abundant in another. For example, various *E. coli* strains spanning a continuum of diversity live in the gut, water, and soil^6^. Analyzing gut metagenomic data alone will most likely reveal a genetic cluster of *E. coli* because it will miss the environmental *E. coli* strains that are either rare or absent in the gut. Unless low abundance strains can be readily recovered or metagenomic sequences from different types of environments are compared, there are also potential pitfalls associated with demonstrating naturally occurring clusters using metagenome-assembled genomes (MAG)^7^. Interestingly, several metagenomic studies have revealed genetic continuum in nature^8,9^.

There is much evidence against the existence of a universal genetic boundary for microbial species. First, the rate of evolution is highly variable across species. Secondly, selection and recombination are thought to be the main cohesive forces driving the formation of genetic clusters. Although recombination rate can be influenced by sequence similarity, there is no correlation between the recombination rate and ANI in bacteria^10^, as recombination can also be affected by physical and ecological barriers. Microbes living in narrow ecological niches and with limited dispersal rate (e.g., obligate intracellular bacteria) may develop genetic clusters. On the other hand, free living microbes exploring different habitats and mixing by dispersal are more likely to exhibit a genetic continuum^11^. Selection is unlikely to produce a universal genetic boundary either, as microbial species are unique in nature, with each species subject to its own evolutionary and ecological forces^12^.

In summary, our study shows that the genetic boundary perceived in the original paper can be explained by persistent within-species sampling bias from historic and current genome sequencing efforts. A more balanced analysis of the present genomic data shows that although genetic clusters may exist in individual species, there is no universal genetic boundary among microbial species in the existing taxonomy.

## Supplementary Materials

### Supplementary Figures

**Supplementary Figure 1.**
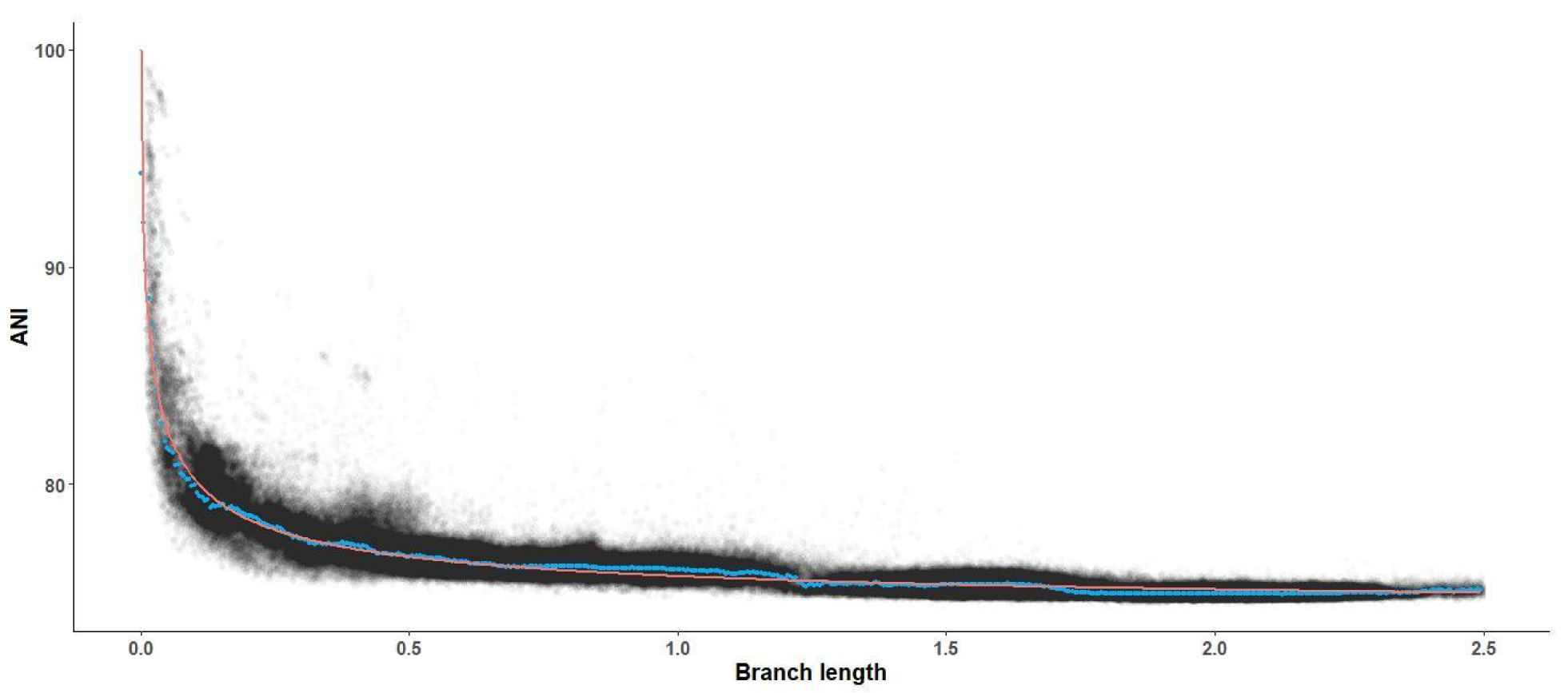
The relationship between the branch length and ANI. Shaded black points are pairwise comparisons between the 3,000 representative genomes. Blue points are the median ANI for each branch length bin (bin width: 0.05 substitution/site). Orange line is the fitted curve. Branch lengths greater than 2.5 were removed because of the lack of data points.

**Supplementary Figure 2.**
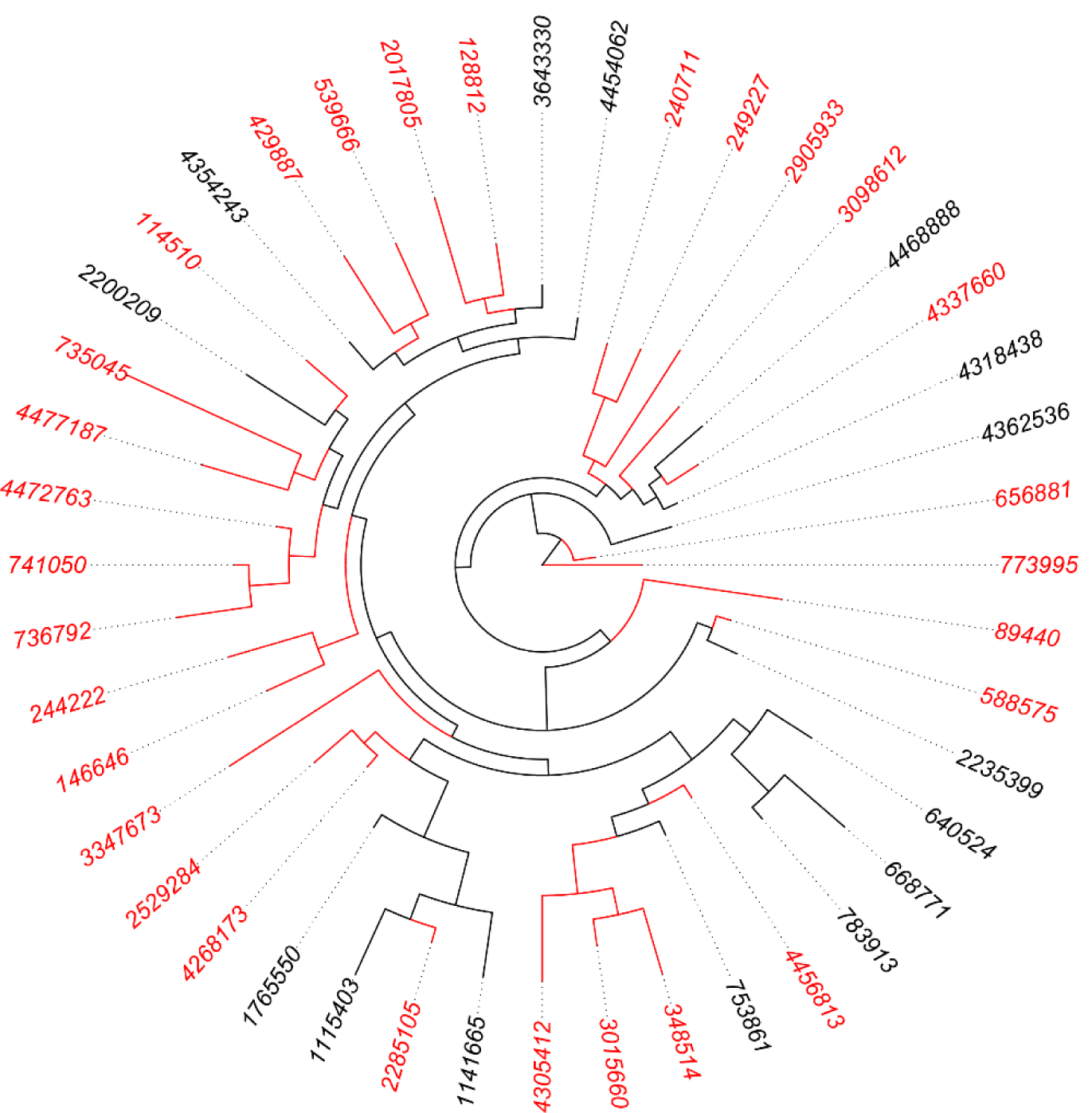
Incomplete and biased coverage of *E. coli* diversity by complete genomes. Of 44 *E. coli* 99% OTUs in the phylogeny of the 16S rRNA gene, only 15 (in black) are represented by the 602 complete *E. coli* genomes in the 10K genome dataset, and only 22% of the total branch length in the phylogeny is covered by these 15 OTUs. The branch length was square-root transformed to better show the short branches in the figure.

**Supplementary Figure 3.**
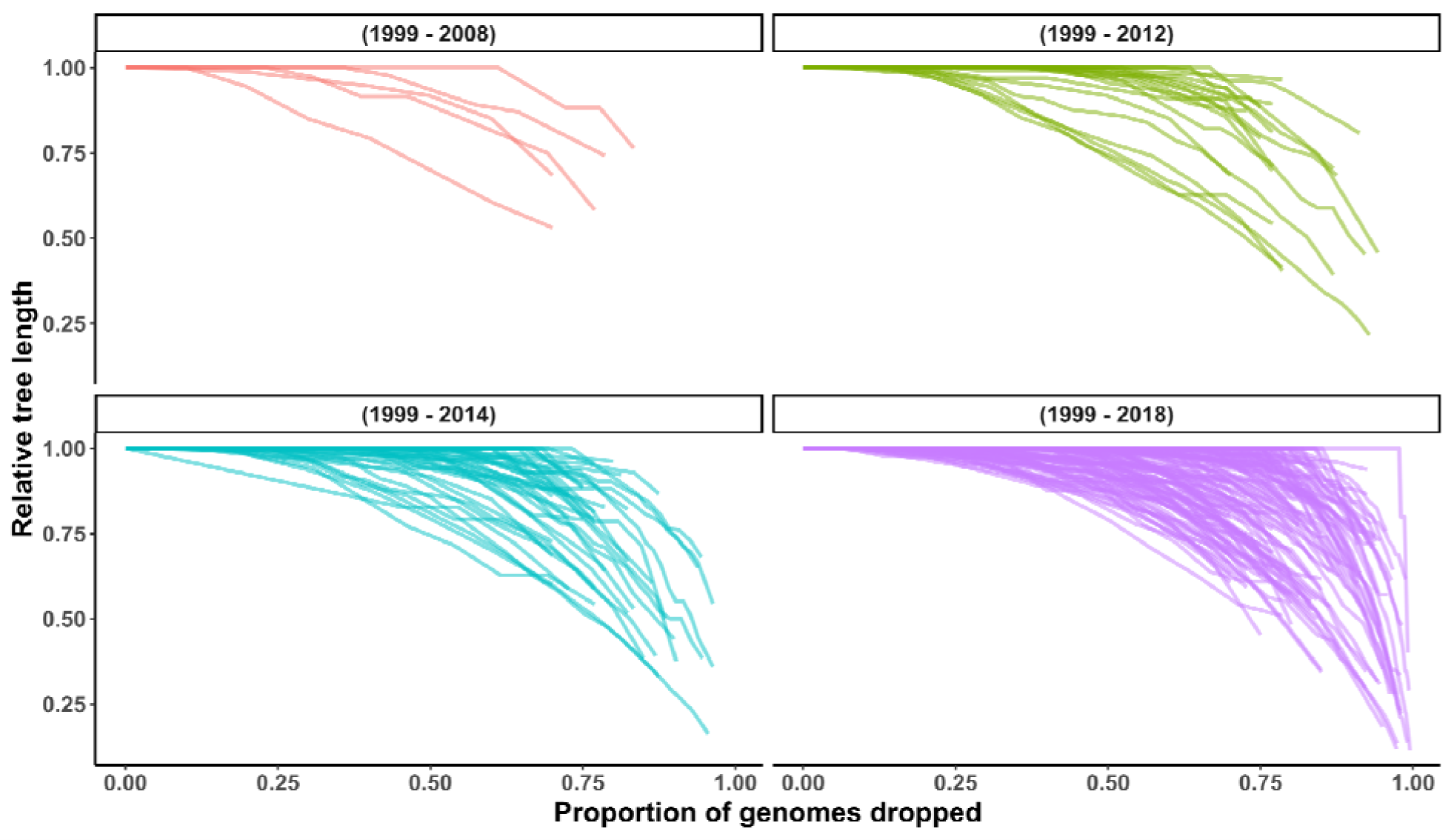
Relative tree length (RTL) decay for species with >= 10 sequenced genomes in the NCBI RefSeq database in four different time periods.

**Supplementary Figure 4.**
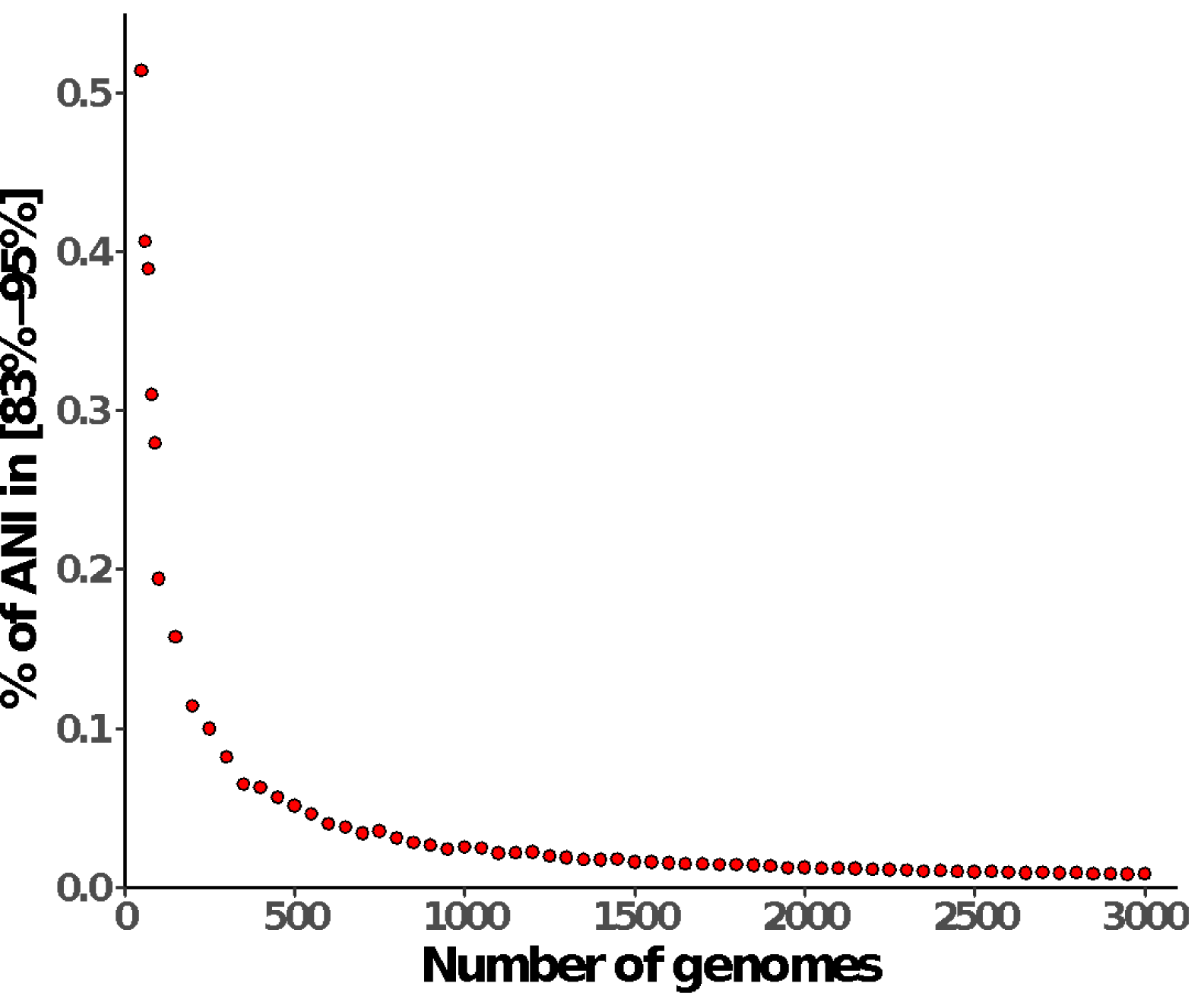
The percentage of ANI values in the [83%-95%] range is marginal and has a negative relationship with the number of genomes. A random phylogeny with an exponential distribution of branch lengths was used to simulate continuous genetic diversity across genomes. The pairwise ANI values were calculated from the phylogenetic distances between genomes (see materials and methods). The simulations were run with phylogenies of different sizes ranging from 50 to 3,000 tips.

## Materials and Methods

### Genome datasets

Two genome datasets were used in this study. The first is the 90K genome dataset from the original paper. It contains both complete and draft bacterial and archaeal genomes. The second dataset consists of 10,616 complete bacterial genomes downloaded from the NCBI RefSeq database on September 6, 2018 (10K genome dataset). From each genome in the 10K dataset, we identified 31 universal protein-coding marker genes using AMPHORA2^13^ and constructed a bacterial genome tree based on the concatenated and trimmed protein sequence alignment of the marker genes using FastTree^14^. Treemmer (version 0.3)^15^ was used to choose 3,000 representative genomes that maximized the phylogenetic diversity in the 10K genome dataset.

### ANI

The ANI values for the 90K genome dataset were downloaded from the original study. For the 10K genome dataset, the 3,000 representative genomes were compared against the full 10K dataset using FastANI (version 1.2).

### Modeling the relationship between branch length and ANI

The 3,000 representative genomes were used to model the empirical relationship between ANI and branch length. The median of ANI was calculated across binned branch lengths (bin width: 0.05 substitution/site) to use as the actual data to fit the relationship between ANI and branch length *l* through the function 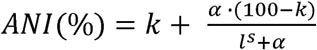, where *k, s* and *α* are shape parameters to be estimated. Minimization of the sum of squares error was performed using the *optim* function in R. The best fit parameters for our data are *α* = 0.075, *k* = 73.94, and *s* = 0.63. Branch lengths greater than 2.5 substitutions/site were removed because of the lack of data points.

### Simulation of continuous genetic diversity and biased within-species sampling

The *rtree* function in the *ape* package in R was used to simulate a random phylogenetic tree of 3,000 tips, with its branch lengths following an exponential distribution with a constant rate of 19.2, estimated from the genome tree of the 3,000 representative genomes. Using the formula described above, the ANI value between a pair of genomes was computed from the branch length between them. To simulate biased sampling within species, a random tip was chosen and two descendants were added to the tip, with the branch length from the tip to the descendant sampled from the same exponential distribution, but its value restricted to the bottom 1% of the distribution. This procedure was repeated on the remaining 2,999 tips until *n* tips were processed this way. Each simulation was run with 10 replicates.

### Assessing the within-species sampling bias

Species with >= 10 genomes were selected from the 10K genome dataset. For each species, a subtree compiling the respective genomes was extracted from the full phylogeny of 10,616 genomes and Treemmer was used to iteratively remove one tip of the tip-pair with the shortest branch length until 3 tips remained. The remaining total branch length of the tree was divided by the total branch length of the initial tree to calculate the relative tree length (RTL) at each iteration. *Rickettsia japonica* and *Chlamydia muridarum* were removed from our analyses because their genomes have identical marker sequences and branch lengths equal to zero.

To further evaluate the within-species sampling bias, we tested how much known genetic diversity is recovered by the complete genomes, using *E. coli* as an example. We extracted 868 unique 16S rRNA gene sequences with no ambiguous bases from 602 complete *E. coli* genomes in the 10K genome dataset and BLAST searched them against 8,655 *E. coli* 16S rRNA gene sequences from the GreenGene 13.8 database. A match was defined as a pair of sequences with 100% identity for their entire sequences. The matched GreenGene 16S rRNA sequences were then mapped to the 99% OTUs (operational taxonomic units) of the GreenGene database. The total branch length covered by the mapped OTUs in the 16S rRNA tree of 44 *E. coli* 99% OTUs was calculated to estimate the coverage of *E. coli* diversity by complete genomes.

### Subsampling of two genomes

For named species with >= 10 genomes in the 90K dataset, two genomes were sampled from each species either randomly or by selecting the pair with the lowest ANI value. Among all pairwise comparisons within the species, the pair with the lowest ANI best represents the phylogenetic diversity of the species.

### Benchmark the performance using 95% ANI for species demarcation

Using the 3,000 representative genomes as the query, we ran FastANI against the full 10K genome dataset. For each query genome, the subject genome with the maximum ANI value was used to benchmark the performance of using the 95% ANI threshold to demarcate bacterial species. A true positive is a query-subject pair belonging to the same species and having an ANI >= 95%. A false positive is a genome pair of different species with an ANI >= 95%, and a false negative is a genome pair of the same species with an ANI of less than 95%. Precision was calculated by: the number of true positive / (number of true positive + number of false positive) and recall was calculated by: the number of true positive / (number of true positive + number of false negative).

## Contributions

M.W. conceived the design of the response, C.S.M., Y.G. and M.W. conducted the analysis, M.W. wrote the first draft and C.S.M., Y.G. and M.W. edited the final version.

